# 50-nm gas-filled protein nanostructures to enable the access of lymphatic cells by ultrasound technologies

**DOI:** 10.1101/2023.06.27.546433

**Authors:** Qionghua Shen, Zongru Li, Matthew D. Meyer, Marc T. De Guzman, Janie C. Lim, Richard R. Bouchard, George J. Lu

## Abstract

Ultrasound imaging and ultrasound-mediated gene and drug delivery are rapidly advancing diagnostic and therapeutic methods; however, their use is often limited by the need of microbubbles, which cannot transverse many biological barriers due to their large size. Here we introduce 50-nm gas-filled protein nanostructures derived from genetically engineered gas vesicles that we referred to as _50nm_GVs. These diamond-shaped nanostructures have hydrodynamic diameters smaller than commercially available 50-nm gold nanoparticles and are, to our knowledge, the smallest stable, free-floating bubbles made to date. _50nm_GVs can be produced in bacteria, purified through centrifugation, and remain stable for months. Interstitially injected _50nm_GVs can extravasate into lymphatic tissues and gain access to critical immune cell populations, and electron microscopy images of lymph node tissues reveal their subcellular location in antigen-presenting cells adjacent to lymphocytes. We anticipate that _50nm_GVs can substantially broaden the range of cells accessible to current ultrasound technologies and may generate applications beyond biomedicine as ultrasmall stable gas-filled nanomaterials.

## INTRODUCTION

Ultrasound has been one of the most widely used imaging modalities in biomedicine, particularly in the fields of obstetrics and cardiology. Yet, the utility of ultrasound can extend well beyond this conventional role of anatomical imaging. Ultrasound-mediated gene/drug delivery, for example, leverages the oscillation of microbubbles caused by an acoustic field to transiently disrupt cell membranes or blood vessel walls for the delivery of genes and drugs^1^. Compared to other *in vivo* gene therapies, such an ultrasound-mediate approach (i) removes the concern of immune responses to viral proteins and the possible vector genome mobilization common; (ii) can noninvasively be focused within most human tissues, enabling the precise control of delivery dose/location; and (iii) is portable, non-ionizing, and broadly available in the clinic. To date, ultrasound-mediated gene/drug delivery has been applied to a wide range of therapeutic cells, including T cells, mesenchymal stem cells, cancer cells, cardiac cells, and cells in the central nerve system^2-9^. Similarly, the use of targeted microbubbles as contrast agents for ultrasound imaging facilitates the acquisition of molecular-level information such as localizing specific biomarkers or cell types^10-12^. Conjugating therapeutic agents to microbubbles also enables the use of ultrasound imaging to monitor the distribution and efficacy of therapeutic agents and potentially, leading to a truly theranostic paradigm^13, 14^.

Unfortunately, microbubbles usually stay within the blood vessels and do not extravasate into tissues due to their relatively large size range between 1-10 μm in diameter (**Fig. 1a**)^15, 16^. As a result, imaging and delivery targets tend to be limited to well-vascularized tissues and can only exert an effect on cells within or very proximal to blood vessels. Although nanobubbles were developed recently to address this challenge, the smallest ones currently still have diameters of ∼ 160 nm^17, 18^, which is too large for crossing many biological barriers. Nanodroplets offer another option of small particles that require acoustic vaporization to become bubbles, though ensuring homogeneity of vaporization *in vivo* and preventing gas loss can be challenging^19, 20^.

**Fig. 1.**
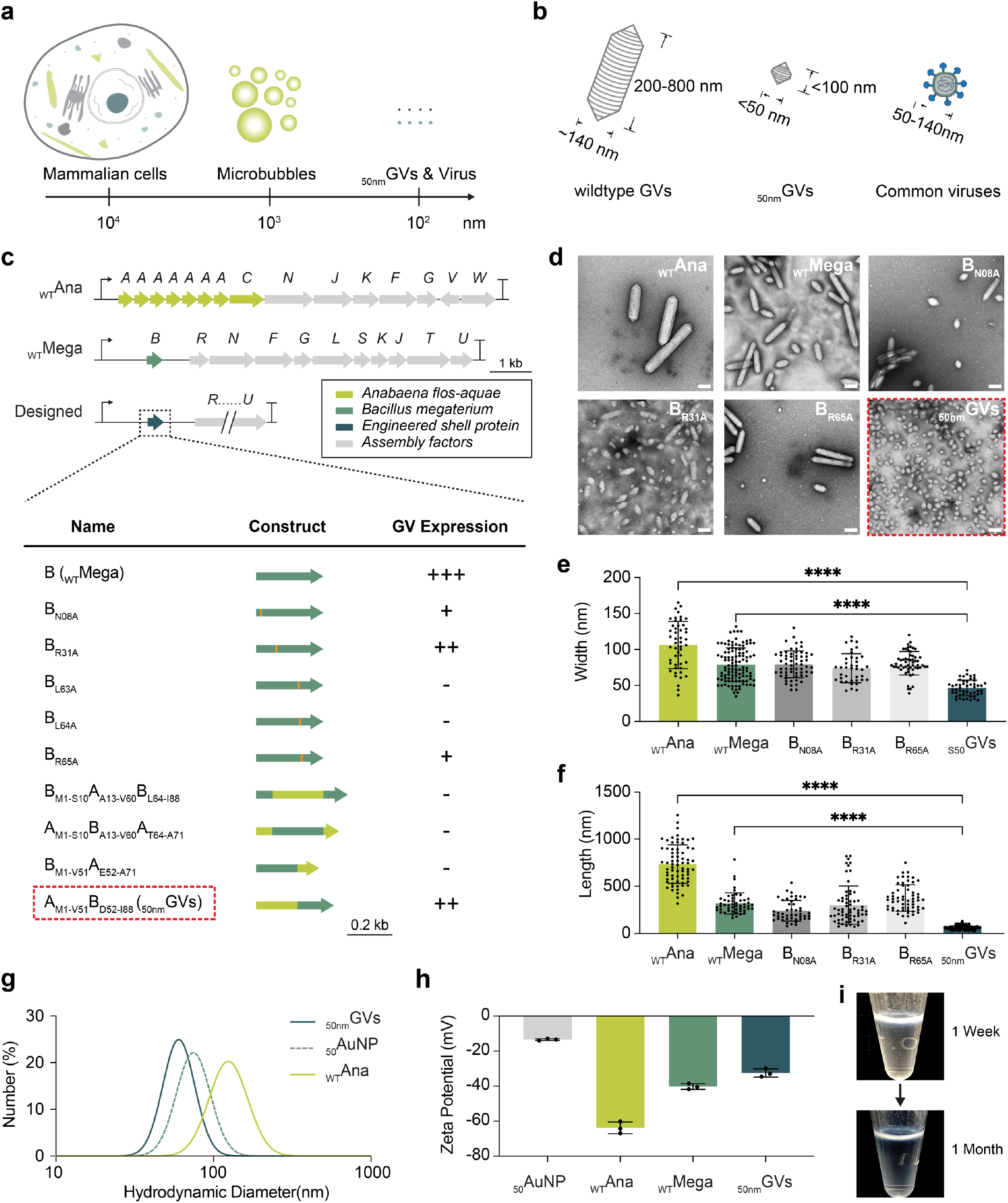
Discovering and characterizing _50nm_GVs. **a**) Schematics of a mammalian cell, microbubbles, _50nm_GVs, and virus particles with their relative scale provided. **b**) Dimensions of wildtype GVs, _50nm_GVs, and common viruses represented by SARS-CoV-2. **c**) Protein sequences of wildtype GVs from *A. flos-aquae* (_WT_Ana), wildtype GVs from *B. megaterium* (_WT_Mega), and engineered GV mutants in this study. Subscripted names stand for mutations, e.g., N08A, or the amino acid segments, e.g., M1-V51. **d**) TEM images of _WT_Ana, _WT_Mega, and _50nm_GVs; scale bar = 200 nm. The final design of _50nm_GVs is highlighted by the red dashed boxes in c) and d). **e, f**) Mean and standard deviation (StdDev) of the particle width and length obtained from TEM images; significance levels: **** p < 0.0001. **g)** Hydrodynamic diameters of _WT_Ana, _50nm_GVs, and _50_AuNPs. **h**) Zeta potential measurements conducted with N = 3 biological replicates. **i**) Representative photos of _50nm_GVs stored in phosphate-buffered saline (PBS) in 1.5 mL tubes at 4°C for 1 week and 1 month, respectively. The visible white layers on the surface of the buffer indicate intact, gas-filled _50nm_GVs.

To gain access to the wide range of cells beyond those near the vasculatures, we hypothesize that it is necessary to develop acoustically active gas-filled nano-agents that are 10-100 nm in diameter. It is well established that the size of nanoparticles plays a crucial role in determining their biodistribution^21^. For example, particles must be 10-100 nm in hydrodynamic diameter to efficiently convect into the lymphatic system^22-24^. Once in, these particles can interact with immune cells and metastatic tumor cells, which play a crucial role in the development of tumor vaccines, early cancer diagnostics, and the treatment of infectious diseases. Similar examples exist such as mucosal tissues, where it is established that < 100 nm particles are required to transport in the mucus and 50-nm virus-like particles may have even better trafficking abilities^25, 26^. Finding out these size limitations is not surprising, as 10-100 nm have been observed to be the common size range of viruses such as SARS-CoV-2 (100 nm), lentivirus (80 nm), and AAV (25 nm), which have evolved to infiltrate biological tissues^27-29^. Consequently, designing ultrasound-responsive nano-agents within this size range is highly desirable as it would significantly extend the reach of current ultrasound technologies to previously inaccessible cell populations.

Here we introduce a new class of 50-nm, stable, free-floating, gas-filled protein nanostructures. They were engineered from the protein sequences encoding gas vesicles (GVs), which are a class of hollow protein nanostructures originally found in photosynthetic microbes^30-32^. Recent biomolecular engineering of GVs opened up multiple applications, including reporter gene imaging by ultrasound, MRI, and optical methods^33-40^, protease sensors^41, 42^, remote cellular control^43, 44^, gas delivery^45-47^, cell tracking^48^, and tumor biomarker imaging^49^. Here we show that genetic mutations of the major shell proteins can shrink both the length and diameter of GVs. Screening of several genetic variants resulted in one that produced a relatively homogenous population of modified GVs with lengths and diameters around 50 nm under transmission electron microscopy (TEM) and hydrodynamic diameters smaller than commercially available 50-nm gold nanoparticles, which we termed as _50nm_GVs. We then focused on the evaluation of the ability of interstitially injected _50nm_GVs to reach deep lymph node tissues and characterized the cell types and sub-cellular compartments they localized to using TEM of ultra-thin tissue sections. The acoustic response of _50nm_GVs was confirmed by the presence of ultrasound imaging contrast. To the best of our current understanding, these _50nm_GVs represent a remarkable achievement as the smallest stable, free-floating bubbles created thus far. Consequently, we envision that our novel agent has potential to extend beyond the realm of biomedicine, opening up possibilities for various applications.

## RESULTS

### The path to discover _50nm_GVs by genetic engineering

GVs were discovered in many phyla of bacteria and archaea. The size of GVs can vary from species to species and is determined by the genetic sequence of the GV operons, which usually consist of ∼ 10 proteins (**Fig. 1b-c**)^30, 50^. Only a handful of GVs have been isolated and characterized to date, and the most commonly observed ones have the shape of a submarine. The center portion of these wildtype GV particles consists of a cylinder, the diameter of which is primarily determined by the sequence of the major shell protein^51^. In contrast, the length of the cylinder is believed to be unregulated, as GVs can continue to grow longer until they reach the physical limit at the boundary of a cell. To reduce the size of GVs to 50 nm, two tasks need to be fulfilled: (1) the diameter of the cylinder needs to be reduced, and (2) the long axis of GVs needs to be shortened. However, the design rules about which amino acids of the major shell protein determine the diameter and length of the cylinder are completely unknown. Thus, the engineering of _50nm_GVs would have to rely on trial and error by testing multiple genetic mutants.

To begin, we chose the pNL29 operon originally cloned from *Bacillus megaterium* (_WT_Mega GVs)^52^. This is because _WT_Mega GVs have the smallest diameter among the commonly studied ones and are amenable to heterologous expression and assembly in *E. coli*, which would allow rapid testing of genetic designs^53^. We first took inspiration from an alanine-scanning mutagenesis study on haloarchaeal GVs^54^, which showed several point mutations that reduced the large diameter of those GVs (mean = 250 nm by negative-staining TEM^53^) into narrow cylindrical ones. Mapping these mutations onto the major shell protein of _WT_Mega GVs, GvpB, resulted in the first set of GV mutants to test: N08A, R31A, L63A, L64A, and R65A (**Fig. 1c-d**). Three of the mutants resulted in successfully assembled GVs. While none of them were able to reduce the diameter of GVs, it was encouraging to see that the N08A and R31A mutants contained a fraction of the shorter, bi-conal GVs.

In a parallel project in the lab^55^, we screened a range of genetic designs that swapped segments of GvpB with another major shell protein from *Anabaena flos-aquae*, GvpA (**Fig. 1c**). From these designs, we uncovered a genetic variant that gave a homogenous population of small bi-conal GVs. Remarkably, the mean diameter of this genetic variant was measured to be 46.4 ± 10.9 nm (mean ± standard deviation) by negative-staining TEM imaging, which is substantially reduced from the ∼ 70 nm mean diameter of _WT_Mega GVs^53, 56^ (**Fig. 1e-f**). Thus, we decided on this GV variant, which consists of the N-terminus to the 2^nd^ β-sheet of *gvpA* from *A. flos-aquae* (residues M1-V51) and the 2^nd^ α-helix to the C-terminus of *gvpB* from *B. megaterium* (residues D52-I88), as the final design of _50nm_GVs.

### Nanoparticle characterization of _50nm_GVs

After establishing the dimensions of _50nm_GVs under TEM, we proceeded to measure the hydrodynamic diameter of _50nm_GVs in a hydrated condition by dynamic light scattering (DLS), which better predicts their behavior in biomedical applications (**Fig. 1g**). To this end, _50nm_GVs showed a hydrodynamic diameter of 63.56 ± 1.26 nm, which was smaller than the 79.21 ± 1.28 nm diameter measured for commercial _50_AuNPs. Notably, having a hydrodynamic diameter smaller than that of the commercial _50_AuNPs is a promising feature, since prior literature has established the favorable *in vivo* biodistribution and application of _50_AuNPs^57^. Next, Zeta potential measurements demonstrated that _50nm_GVs have negatively charged surfaces similar to _WT_Ana and _WT_Mega GVs (**Fig. 1h**). Lastly, we used the dimensions of _WT_Ana, _WT_Mega, and _50nm_GVs measured from negative-staining TEM images to estimate the molecular weight and gas volume of _50nm_GVs (**Table 1**) as previously described^53^. This calculation revealed that the molecular weight of _50nm_GVs is ∼ 10 times and the volume ∼ 25 times smaller than those of _WT_Mega GVs.

**Table 1.** Properties of gas vesicle variants.

| GV variants: | $_{WT}Ana$ | $_{WT}Mega$ | $_{50nm}GVs$ |
| --- | --- | --- | --- |
| Diameter, mean value (nm) | 106.1 | 78.9 | 46.4 |
| Diameter, standard deviation (nm) | 32.9 | 23.3 | 10.9 |
| Length, mean value (nm) | 734.6 | 319.5 | 63.2 |
| Length, standard deviation (nm) | 203.2 | 111.3 | 21.1 |
| Estimated GV molecular weight (MDa) | 252.57 | 77.17 | 7.71 |
| Estimated GV volume (attoliter) | 4.07 | 0.850 | 0.0293 |
| Estimated gas volume (attoliter) | 3.77 | 0.759 | 0.0202 |
| Est. gas vol. per mg of protein ($\mu L/mg$) | 8.98 | 5.92 | 1.58 |

### _50nm_GVs are fundamentally stable “bubbles”

Another key distinction of GVs from synthetic microbubbles is that GVs are fundamentally stable, owing to a design principle that allows air exchange but prevents water condensation by the inner hydrophobic surface^50, 58^. The extended stability of _50nm_GVs presents a distinct advantage for various applications, including the targeting of lymph nodes, as described herein, given that trafficking through interstitial spaces and the lymphatic system is often a protracted process spanning several hours (see the next section). Another example that requires long-term stability of the gas-filled nano-agents is multi-day tracking of mesenchymal stem cells, as done in a recent study^48^. Here we validated the stability of _50nm_GVs by monitoring a tube of _50nm_GVs during storage for over a month and confirming that the floating white layer remained stable, supporting that _50nm_GVs have a similar stability profile as other wildtype GVs (**Fig. 1i**).

### _50nm_GVs can penetrate the barrier of lymphatic endothelial cells and gain access to immune cells

To evaluate the ability of _50nm_GVs to reach previously inaccessible cell populations, we focused our study on the biodistribution of _50nm_GVs in lymph nodes. To this end, lymphatic endothelial cells (LECs) present one of the most heavily studied biological barriers due to their importance in determining the delivery efficiency of therapeutics to immune cells in the lymph node. It has been previously shown that only nanoparticles 10–100 nm in hydrodynamic radius can efficiently transverse lymphatic endothelial cells into lymphatic tissues^22-24^. Experimentally, we first conjugated Alexa fluorophores to all 3 types of nanoparticles in our study, _50nm_GVs, _WT_Ana GVs, and _50_AuNPs, before interstitially injecting an approximately equal amount of each type into the front paw of mice (**Fig. 2a**). The trajectory of particle trafficking through the interstitial space to the adjacent axillary lymph node was tracked using fluorescence live animal imaging. The findings from the study indicate that the two smaller particles, _50nm_GVs and _50_AuNPs, reached the axillary lymph node region approximately 60 minutes after injection. In contrast, the larger _WT_Ana GVs took around 90 minutes to reach the same location, which aligns with the expected transport behavior due to lymphatic drainage (**Fig. 2b-c**).

**Fig. 2.**
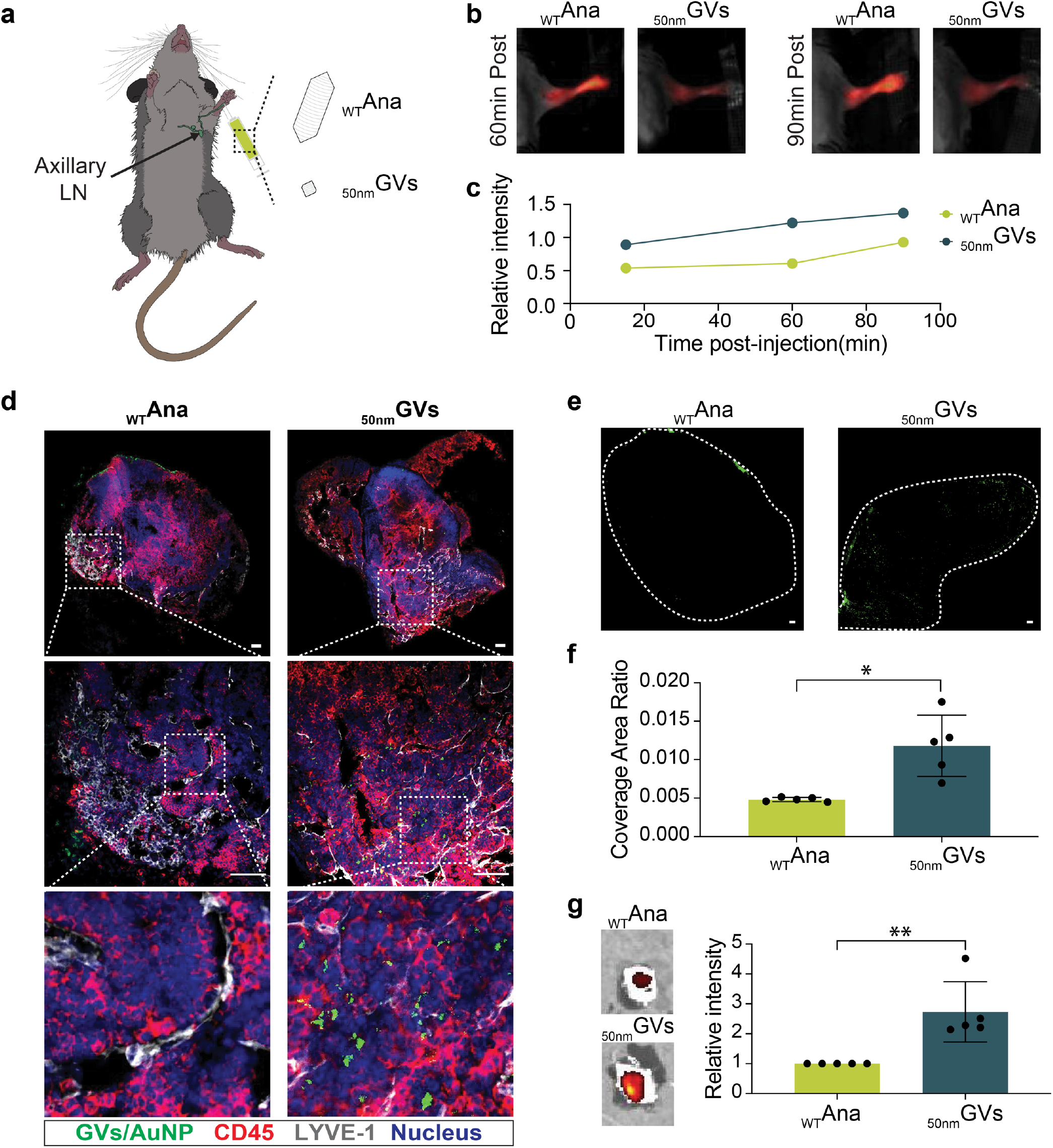
_50nm_GVs can penetrate the barrier of lymphatic endothelial cells and gain access to immune cells. **a**) Schematic representation of the injection site and the targeted lymph node. **b**) *In vivo* imaging system (IVIS) images showing the transportation kinetics of injected _WT_Ana and _50nm_GVs. **c**) Quantitative analysis of the change in fluorescent intensity within the targeted lymph node area at different timepoints (30min, 60min, and 90min post-injection). Relative intensity is calculated as lymph node area intensity divided by injection site intensity. **d)** Confocal fluorescence images of immunohistochemistry to depict the distribution of _WT_Ana and _50nm_GVs within the lymph node tissues. The white dashed boxes in the top row images outline the relative location of the zoomed-in areas depicted in the second row, and the white dashed boxes in the second row indicate the further zoomed-in areas, which are shown in the bottom row. Scale bar = 50 μm. The images are an overlay of three images acquired in the red fluorescence channel that showed expression of CD45, the green fluorescence channel that showed the location of the nanoparticles, and the blue fluorescence channel that showed expression of LYVE-1. **e**) Visualization of particle distribution within the dissected lymph nodes with green-fluorescent dots representing injected particles and white dashed lines outlining the periphery of each lymph node. Scale bar = 50 μm. **f**) Quantification of the total coverage area of particles inside lymphatic tissue. Results are presented as the ratio of particle area to the entire lymph node area. Significance levels: * p < 0.05. **g)** Total fluorescent intensity of lymph nodes dissected from mice injected with _WT_Ana and _50nm_GVs. The Y-axis represents the _50nm_GVs:_WT_Ana fluorescent intensity ratio of each experimental pair with samples prepared in parallel. Significance levels: ** p < 0.01. See additional datasets in Fig. S2 and Fig. S3.

We then euthanized the mice and dissected the lymph nodes for immunohistological analysis (**Fig. 2d**). Antibodies against lymphatic endothelial cells surface receptor LYVE-1 were used to label lymphatic vessels, and antibodies against CD45 were to label the lymphocytes. To determine if a type of nanoparticles has successfully infiltrated into lymphatic tissues, we should focus on regions that have CD45 staining and are in a certain distance away from lymphatic vessels labeled by anti-LYVE-1 antibodies. Indeed, _50nm_GVs were observed in discrete puncta co-localized with the CD45 staining and situated apart from the LYVE-1 staining. Conversely, _WT_Ana GVs were detected in a limited quantity solely on the outer surface of the lymph nodes, indicating a distribution within the capsular and subcapsular spaces of lymph nodes. As a positive control, _50_AuNPs showed a similar distribution pattern as _50nm_GVs, agreeing with previously reported biodistributions of ∼50-nm nanoparticles^59^ (**Fig. S3**). To quantify the biodistribution, we analyzed the total areas that the two types of GVs covered on tissue slides, and the results showed that _50nm_GVs covered areas that are nearly twice larger on average than that of _WT_Ana (**Fig. 2e-f**). The image analysis was further corroborated by a measurement of total fluorescent intensity of dissected lymph nodes, which revealed consistently higher fluorescent intensity for the _50nm_GVs group compared to the _WT_Ana GVs group (**Fig. 2g**). Together, these data support that _50nm_GVs are able to extravasate from lymphatic vessels and potentially access lymphocytes, as predicted from the previously reported size threshold for nanoparticles to transverse through lymphatic endothelial cells^22^.

### Cell-type and subcellular distribution of _50nm_GVs revealed by TEM images

Next, we acquired TEM images of ultra-thin tissue sections to examine and compare the cell-type and subcellular distributions of _WT_Ana GVs and _50nm_GVs. First, we screened through different lymph node regions, from their outer surface to deeper-lying tissues (**Fig. 3a-b**). For the lymph nodes exposed to _WT_Ana GVs, only a limited amount of GVs were observed within the subcapsular sinus, agreeing with the immunohistological images. Also, these GVs appeared to be shorter than the average length expected for _WT_Ana GVs, indicating that capsular structures of lymph nodes generally prohibit the transport of the longer wildtype GVs into lymph nodes (**Fig. 3c**). In contrast, _50nm_GVs were observed both in the subcapsular area and deeper lymphatic tissues (**Fig. 3d-e**). In most of the cases, _50nm_GVs were observed in large cohorts within the endosomal compartments of antigen-presenting cells, such as the phagosomes of dendritic cells and macrophages (**Fig. 3f**). Importantly, cells that carried these _50nm_GV-containing compartments were observed to be immediately adjacent to lymphocytes in multiple instances (**Fig. 3g**), supporting the potential use of _50nm_GVs to deliver payloads to these antigen-presenting cells that may be in direct contacts with lymphocytes. We also observed _50nm_GVs in compartments near the subcapsular sinus (**Fig. 3h**), encouraging further analysis of their trafficking routes. Lastly, clusters of _50nm_GVs were identified within lymphatic vessels, suggesting their predominant delivery route occurs via the lymphatic drainage system (**Fig. 3i**). Overall, the higher-resolution tissue images provided by TEM further confirmed the delivery of _50nm_GVs to the deep lymphatic tissues, and the large amount of _50nm_GVs that reached the vicinity of lymphocytes paves the way for future engineering of _50nm_GVs with specific payloads and targeting moieties.

**Fig. 3.**
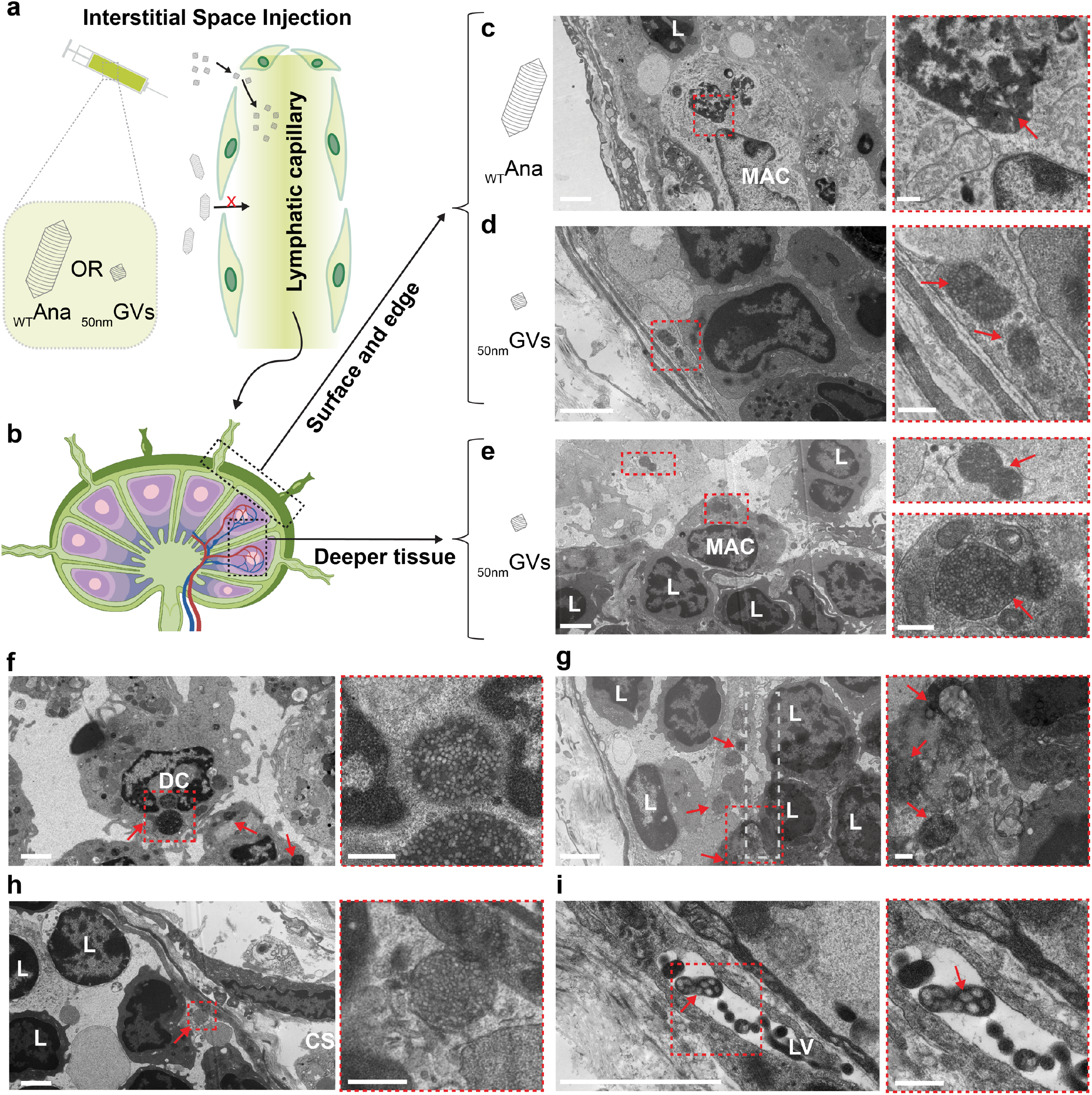
Cell-type and subcellular distribution of _50nm_GVs inside lymphatic tissues revealed by TEM images. **a**) Schematic representation of the lymphatic tissue barrier and nanoparticles pathway after interstitial injection. **b**) Anatomical schematic of a lymph node with two windows that indicate different depths into the tissue. The black arrow indicates the relevant TEM image collected from that area. **c**) Representative TEM images of _WT_Ana GVs observed in the outer surface regions of the lymph nodes corresponding to capsular and possible subcapsular sinus. Red arrows indicate the location of GVs. **d-e**) Representative TEM images of _50nm_GVs observed in the outer surface and deeper tissue regions. Red arrows indicate the location of _50nm_GVs in endosomal compartments of macrophages that are often surrounded by lymphocytes in lymphatic tissues. See also Fig. S4 for additional detail on cell type identification in TEM images. **f)** A representative image of a dendritic cell with _50nm_GV-containing phagosomes. **g**) A representative image of _50nm_GVs in compartments immediately adjacent to lymphocytes. A white dashed box indicates possible contact areas between the antigen-presenting cells and lymphocytes. **h**) A representative image of _50nm_GVs in compartments immediately adjacent to the capsular sinus. **i**) A representative image of _50nm_GVs in small clusters inside lymphatic vessels. The abbreviations used are CS for capsular sinus, L for lymphocytes, LV for lymph vessels, MAC for macrophages, and DC for dendritic cells. (b) is created with BioRender.com. For c-i, the right images with a red dashed outline indicate the zoom-in areas from the labeled regions in the corresponding left images. Scale bars are 2 μm for the left images and 400 nm for the right images in each panel.

### _50nm_GVs are acoustically active

While the fact that _50nm_GVs can be buoyancy-purified proves that they are gas-filled, we sought to further confirm that the gas compartments of _50nm_GVs are acoustically active in a similar fashion to their wildtype counterparts, especially at ultrasonic frequencies. To this end, we carried out ultrasound imaging and measured the signal generated by _50nm_GVs. Inspired by the recently developed BURST method that leveraged the collapse of GVs under high ultrasound pressure to achieve higher sensitivity^35^, we mimicked the BURST imaging (i.e., focused transmits) instead of plane-wave imaging to ensure the reliable collapse of _50nm_GVs by a 21-MHz transducer (**Fig. 4a**). Experimentally, a series of frames were recorded at higher ultrasound transmit pressures to capture the collapse process, and then the max intensity projections of the second half of the series (*i*.*e*., post-collapse) were subtracted from the first half (*i*.*e*., pre-collapse) to give the final ‘BURST’ images (**Fig. 4b**). Under this scheme, collapsible ultrasound contrast agents present significant contrast in the subtraction image, while normal biological tissues and non-collapsible agents such as polystyrene beads, do not. As expected, when we loaded _50nm_GVs, _WT_Ana GVs, and polystyrene beads (_Ctrl_PS) into an agarose phantom, we observed strong ‘BURST’ signals from both types of GVs, while _Ctrl_PS expectedly failed to provide contrast in the ‘BURST’ image (**Fig. 4c**). Plotting average image intensity as a function of frame number further revealed a ‘BURST’ peak for both _50nm_GVs and _WT_Ana GVs (**Fig. 4d)**. Next, we measured the ultrasound signal intensity from serial dilutions of _50nm_GVs and _WT_Ana GVs. Both agents presented with concentration-dependent signals (**Fig. 4e**). At the same protein concentration, _50nm_GVs produced weaker signals than _WT_Ana GVs, which is expected due to their lower gas-to-protein ratio (**Fig. 4f** and **Table 1**). In conclusion, these results establish that _50nm_GVs are responsive to ultrasound, similar to their wildtype counterparts. The measured ultrasound signal intensity of _50nm_GVs in relation to _WT_Ana GVs will serve as a guideline for their usage in contrast-enhanced ultrasound imaging and molecular imaging of biomarkers.

**Fig. 4.**
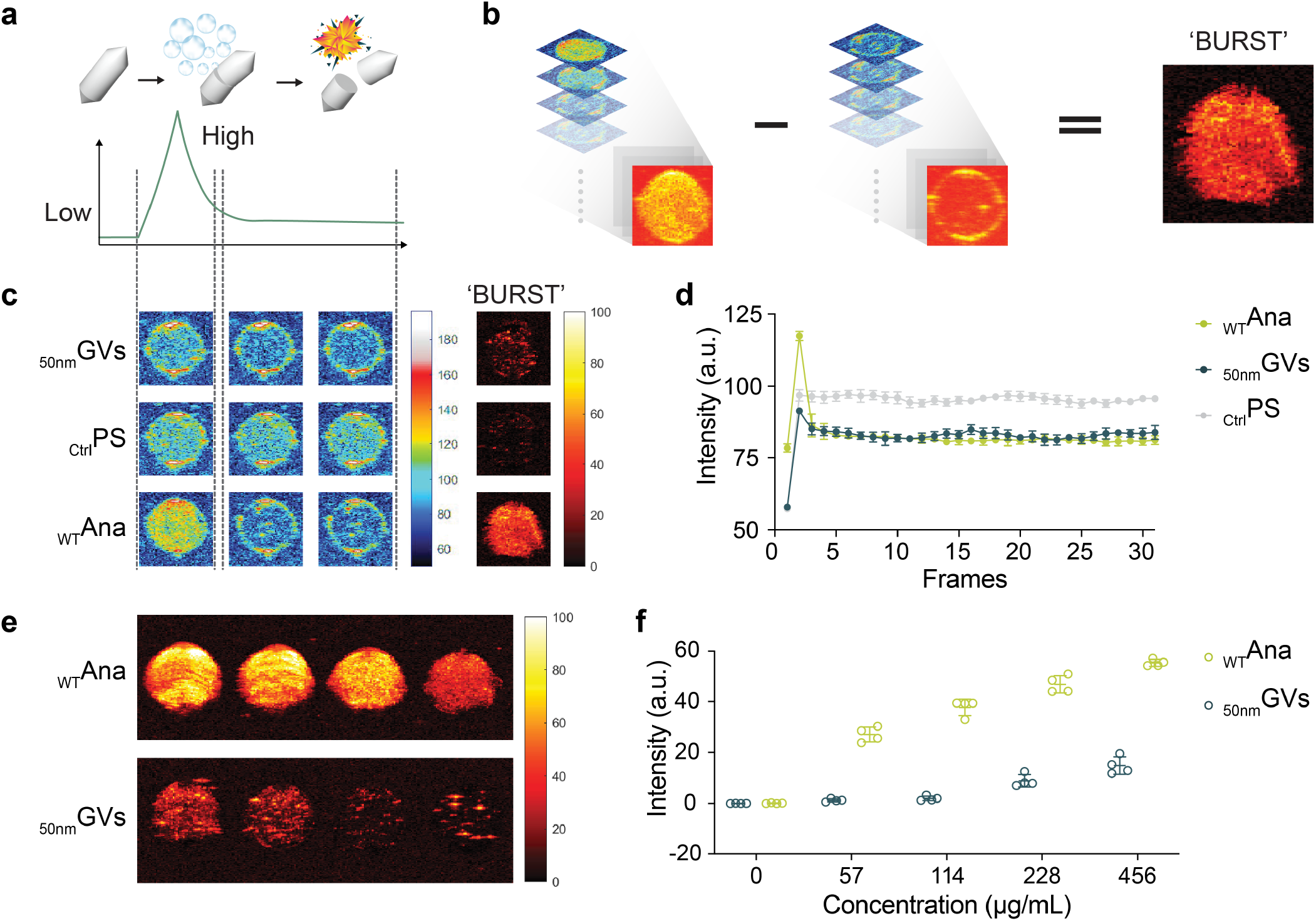
_50nm_GVs are acoustically active. **a**) Schematic diagram illustrating the collapse process of GVs. The light green line indicates the acoustic pressure, and the dark green line indicates the ultrasound signal from GVs. **b**) Schematic diagram demonstrating the process of generating a final ‘BURST’ image by subtracting B-mode images taken from the earlier frames from those from later frames. **c**) Representative ultrasound images of the B-mode images (1^st^, 3^rd^, and 30^th^ frames) during the high-pressure collapse process and the final ‘BURST’ image. _Ctrl_PS refers to polystyrene beads used as a control sample. **d**) Average intensity within N = 4 representative regions of interest (ROIs) for GVs and _Ctrl_PS as function of frame number. **e**) BURST image of a phantom containing serial dilutions of _WT_Ana and _50nm_GVs with protein concentrations of 456 μg/mL, 228 μg/mL, 114 μg/mL, and 57 μg/mL, respectively. **f**) Quantification of the signal intensity target ROIs from images in (e) and other replicates. Error bars representing mean ± standard deviation for N = 4 replicates.

## DISCUSSION

The _50nm_GVs described in this study, to the best of our knowledge, appear to be the smallest stable, free-floating bubbles created to date. In contrast, gases within microbubbles, nanobubbles, and those vaporized from nanodroplets typically exist in a non-equilibrium state. Although 1-nm bubbles have been described in material science, their requirement for a graphene interface for formation renders them non-free-floating^60^. Therefore, the development of _50nm_GVs holds the potential to have an impact not only in biomedicine but also in areas such as material synthesis and the exploration of interfacial chemistry.

Furthermore, this study represents a pioneering report on the biodistribution of various genotypes of GVs within lymph nodes. Since lymph nodes have been a focal spot for immunological studies and therapeutic developments such as cancer vaccines^61^, our study may constitute the first step toward applying ultrasound-based imaging and delivery methods to these previously inaccessible lymphatic cells.

Given the ultrasmall dimensions of _50nm_GVs, we postulate that their size may facilitate their traversal across diverse biological barriers, extending beyond penetration of lymphatic endothelial cells, as demonstrated in our study, to perhaps also include penetration of the blood-brain barrier (BBB) ^62^ as previous studies indicate nanoparticles between 50-200 nm would be ideal to cross this interface^63, 64^. These findings hold promising implications for potential applications of _50nm_GVs in treatment of neurological disease.

Lastly, it will be important to formulate design rules for ultrasmall GVs and pinpoint the protein sequences that encode their structural features. While 8 proteins were identified to be essential for the assembly of GVs^34^, there is currently little understanding as to how these proteins work together at a molecular level. This presents a key bottleneck in decoding the genetic design rules for ultrasmall GVs. We anticipate that our study here, together with a companion paper by Lin *et al*. that presents another type of ultrasmall GVs with a different genetic sequence, will provide the initial genetic sequences to spur studies on their protein structures, opening the door to a fuller understanding of the molecular mechanism behind the remarkable formation of these ultrasmall, stable, free-floating bubbles.

## MATERIALS AND METHODS

### Cloning, expression, and purification of GVs

Plasmids pST39-pNL29 and ARG1 were obtained from Addgene to encode GV gene clusters from *B. megaterium* and *gvpA* from *A. flos-aquae*, respectively (#91696 and #106473), and from these two plasmids, we constructed the genetic sequence of the major shell protein of _50nm_GVs using Q5 DNA polymerase and Gibson assembly kits (New England Biolabs Inc.)^65^. To express _50nm_GVs, point mutant GVs (N08A, R31A, L63A, L64A, and R65A), and _WT_Mega GVs, we transformed the plasmids into BL21 Star™ (DE3) pLysS One Shot™ *E. coli* and cultured them in LB Miller Broth (Thermo Fisher Scientific, Waltham, MA) with 100 μg/mL Carbenicillin (Gold Biotechnology, Olivette, MO), 25 μg/mL Chloramphenicol (MilliporeSigma, Burlington, MA), and 0.2% glucose (MilliporeSigma, Burlington, MA). When the cells reached an OD_600_ of 0.6, 80 μM (for _50nm_GVs) or 20 μM (for all others) Isopropyl β-d-1-thiogalactopyranoside (IPTG) (Teknova, Hollister, CA) was added to induce the protein expression for 22 hrs at 28.5 °C. Cells were collected and pelleted by centrifugation at 400 x g in 50 mL conical tubes, and then the middle layer was removed to isolate the remaining cells. The cells were mixed with SoluLyse-Tris, lysozyme (Genlantis, San Diego, CA), and DNase I (MilliporeSigma, Burlington, MA), and the lysate was transferred to 2 mL tubes for three cycles of centrifugally assisted flotation to further purify the GVs. To uncluster the _50nm_GVs and _WT_Mega GVs, GVs in phosphate-buffered saline (PBS) were added with urea to a final concentration of 6 M, followed by gentle agitation at room temperature for 1 hr as previously described^53^. After this period, the GV solution was resuspended in 1× PBS and centrifuged at 350 × g overnight at 4°C. This process was repeated 3 times to ensure the complete removal of bottom urea media. Expectedly, this process results in a substantial reduction in the optical density at 500 nm (OD_500_) of the GVs and thus the OD_500_ of _50nm_GVs referenced in the paper was measured after the unclustering procedure. To produce _WT_Ana GVs, *A. flos-aquae* (CCAP strain 1403/13 F) was cultured and harvested as previously described^53^. The floating cells were lysed using sorbitol and Solulyse solution (Genlantis), and GVs were separated from debris through repeated centrifugally assisted floatation.

### Hydrodynamic size measurement

To measure the hydrodynamic diameter of clustered and un-clustered GVs, purified GV samples were diluted to OD_500_ = 0.2 in 1X PBS, and 500 μL of each sample was transferred into a cuvette (Thermo Fisher Scientific, Waltham, MA). The measurements were taken using a Malvern Zen 3600 Zetasizer (Malvern, UK) with a minimum of three measurements per sample. At least three biological replicates were conducted for each type of GV.

### Interstitial injection and fluorescence live animal imaging

To monitor the migration of GVs to the lymphatic system, GVs (230 μg/mL), and 50 nm amine_gold nanoparticles (Cytodiagnostics, Canada) were first labeled with NHS-Alexa488/694 (Thermo Fisher Scientific, Waltham, MA). These fluorescently labeled GVs, and gold nanoparticles were then dissolved in 20 μL of PBS under sterilized conditions. Following previously described procedures^66-68^, prepared solutions were injected directly into the interstitial space of the BALB/c mouse paw (N = 3 per group) to allow the uptake of the nanoparticles into the lymphatic system. To track the nanoparticle signals after the injection, the animals were imaged at 60 and 90 minutes post-injection using IVIS Lumina II (Advanced Molecular Vision) following the manufacturer’s recommended procedures and settings. To analyze lymph node accumulation, the intensity of the axillary lymph node was measured using a region of interest (ROI) within the representative area. The lymph node’s relative intensity was calculated as the ratio of the lymph node intensity to the primary injection site intensity, i.e., relative intensity = (lymph node intensity) / (primary injection site intensity). To collect samples for immunohistology, mice injected with _50nm_GVs were euthanized at 60 minutes post-injection, and _WT_Ana GVs, at 90 minutes post-injection. In all cases, the targeted axillary lymph node reached a similar relative intensity (∼0.9). Axillary lymph nodes were then dissected for immunostaining following standard staining protocols. All animal procedures were approved by the Institutional Animal Care and Use Committee (IACUC) of Rice University and performed according to the guidelines.

### Immunohistology

To further analyze the dissected lymph node tissues from all experimental groups, including _50nm_GVs, _WT_Ana GVs, and _50_AuNP, the lymph node tissues were fixed with 4% paraformaldehyde and sectioned into 15 μm slides. Immunohistochemistry was performed to evaluate the expression of specific markers, LYVE-1 and CD45. The slides were first blocked for 1 hour with 5% goat serum (MP Biomedicals, California) in a washing solution consisting of 0.5% Triton X-100 in 1x PBS. The primary antibodies used in this study were LYVE-1 (E3L3V) Rabbit mAb (cat. no. 67538S, Cell Signaling, Danvers, MA) and CD45 (30-F11) Rat mAb (cat. no. 55307S, Cell Signaling, Danvers, MA). The antibodies were diluted 1:200 in 5% goat serum and incubated overnight at 4°C. Following incubation, the slides were washed three times with the washing solution for 5 minutes each. The secondary antibodies, Anti-rat IgG (H+L), Alexa Fluor 647 (cat. no. 4418, Cell Signaling, Danvers, MA), and Anti-rabbit IgG (H+L) Alexa Fluor 350 (cat. no. 11046, Invitrogen, Waltham, Massachusetts) were diluted 1:200 in 1 x PBS. The slides were then incubated with the secondary antibodies for 1 hour at room temperature. The images were acquired using a Nikon A1R-si Laser Scanning Confocal Microscope (Japan), equipped with a laser of 405/488/561/638 nm. The expression of LYVE-1 and CD45 in the tissue was evaluated by examining the staining pattern of each antibody within the lymph node tissue samples.

### Transmission electron microscopy

To prepare the lymph tissue samples for TEM, extracted tissue was fixed overnight at room temperature in Karnovsky’s fixative (Electron Microscopy Sciences, Hatfield, PA) and then post-fixed for one hour in 1% osmium tetroxide. Samples were dehydrated in a graded series of ethanol, embedded in epoxy resin and polymerized overnight at 70 °C. Ultra-thin sections of 100 nm thickness were cut using an ultramicrotome (Leica EM UC7), placed on an unsupported 200-mesh copper grid, and then post-stained with saturated methanolic uranyl acetate and Reynold’s lead citrate. Images were collected using a JEOL JEM-1400Flash TEM operating at 120 kV and equipped with an AMT NanoSprint15 sCMOS sensor. For TEM of negatively stained purified GVs, samples were diluted to OD_500_ = 0.2 in 1 X PBS and loaded onto 200-mesh carbon-coated copper grids (Ted Pella, Redding, CA) for three minutes. Excess liquid was carefully blotted away with filter paper, and the samples were then stained with 2% (w/v) uranyl acetate (Electron Microscopy Sciences, Hatfield, PA). High-resolution TEM images were captured using a JEOL JEM-2010 TEM and a JEOL JEM-2100F TEM.

### Ultrasound imaging

Imaging phantoms were fabricated by preparing a 1% agarose solution in PBS. Various concentrations of GVs in PBS were mixed with a PBS solution containing 2% agarose in a 1:1 ratio at 50°C, immediately followed by loading 150 μL of the resultant mixture into wells in the phantom. Imaging was performed using a Vevo F2 system (FUJIFILM Visualsonics Inc., Toronto, Canada), equipped with a linear array (UHF29x) with a 21-MHz center frequency. For transmit pulses, the center frequency is 21 MHz, the f-number is 2.0, and the number of cycles is 1. All images were taken with the VADA interface using customized sequences and processed using MATLAB. B-mode BURST sequences consist of a single low-pressure frame (11% output power at the VADA interface), followed by 30 repetitions of 5 high-pressure (75% output power) frames with different focal depths between 9 and 13 cm to cover the entire sample-containing region. Thus, a total of 151 frames were acquired in each series. The 5 frames of different focal points were first averaged, and the resulting 30 frames were split into two halves. For each half, a single frame was formed using pixel-wise maximum intensity projection of the 15 frames, and the final BURST image was generated by pixel-wise subtraction of the second frame from the first one.

### Quantification and statistical analysis

Information on sample size (n) and P-values for the experiments can be found in the figures, figure captions, and method section. Statistical analysis was performed using GraphPad Prism software and presented as mean ± standard deviation (StdDev). Independent t-test were conducted for the comparison between two groups. Multiple comparisons were analyzed using Welch and Brown-Forsythe ANOVA tests, which do not assume equal variances across all groups in a population. Significance was set at p<0.05.

## ACKNOWLEDGMENTS

We thank the Shared Equipment Authority (SEA) at Rice University for the access to core facilities and instruments, Dr. Wenhua Guo for the training, and Drs. Melissa Yin and Isabel Newsome from FUJIFILM Visualsonics Inc. for technical support. This work was supported by the Cancer Prevention and Research Institute of Texas (CPRIT), the NIH (R00 EB024600 and R21 EB033607), the Welch Foundation, the G. Harold and Leila Y. Mathers Foundation, the Hearing Health Foundation, and the John S. Dunn Foundation.

## AUTHOR CONTRIBUTIONS

Conceptualization, G.J.L.; Methodology, G.J.L., Q.S., Z.L., R.R.B. and M.D.M.; Investigation, Q.S., Z.L., M.D.M., M.T.D., and J.C.L.; Formal Analysis, Q.S., Z.L., M.D.M., M.T.D., and J.C.L.; Writing – Original Draft, G.J.L., Q.S., Z.L., and R.R.B.; Supervision and Funding Acquisition, G.J.L.

