## Supplemental materials for "50-nm gas-filled protein nanostructures to enable the access of lymphatic cells by ultrasound technologies"

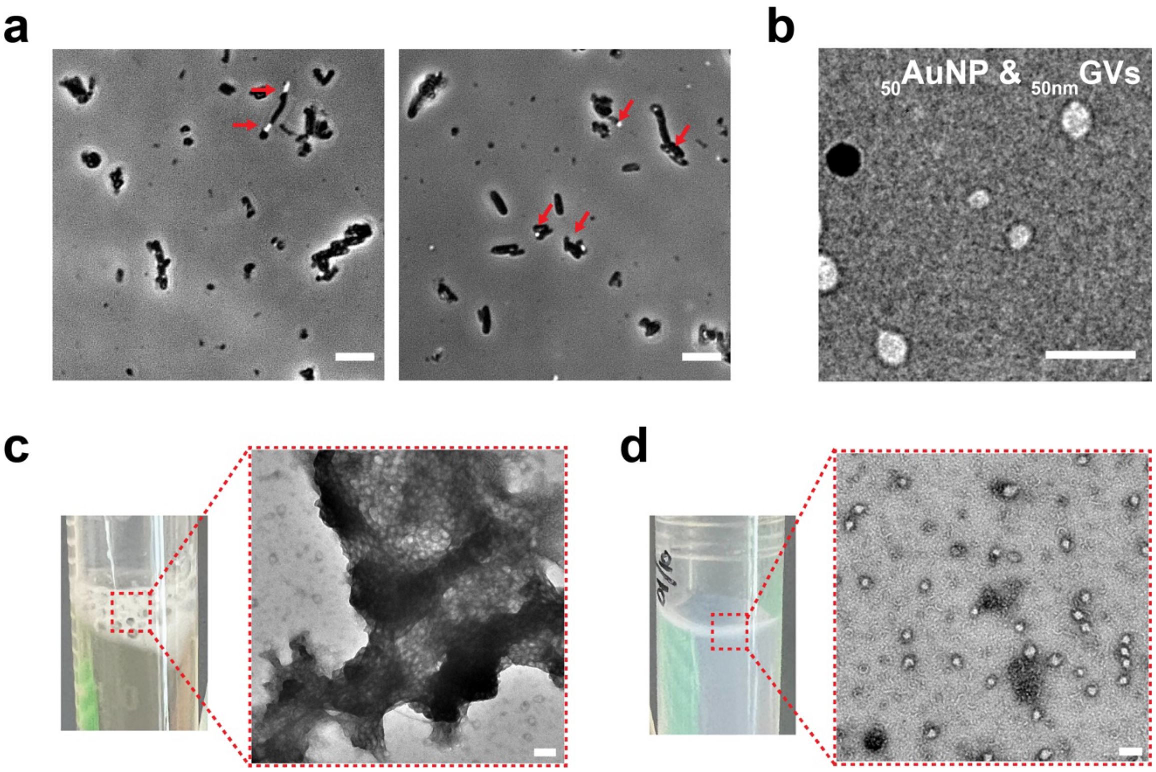


**Figure S1. Purification and additional characterization of $_{50nm}$GVs.** a) Representative phase contrast images of $_{50nm}$GV-expressing *E.coli*. *E.coli* cells with strong GV expression have visible reflecting dots in the phase contrast imaging (Farhadi et al., 2020) as indicated by the red arrows. Scale bars = 5 µm. b) A transmission electron microscopy (TEM) image of $_{50nm}$GVs mixed with commercial 50-nm gold nanoparticles ($_{50}$AuNPs) for the comparison of their structural properties. $_{50}$AuNPs appear as darker contrast and GVs appear as lighter contrast on TEM images. Scale bar = 200nm. (c) A representative camera image of a 1.5 mL tube containing floating $_{50nm}$GVs and a TEM image of clustered $_{50nm}$GVs. Right after cell lysis, the clustered $_{50nm}$GVs are visually discernible as a floating white layer. d) A representative camera image and TEM image of $_{50nm}$GVs unclustered by 6 M urea treatment as described in previous protocols (Lakshmanan et al., 2017).

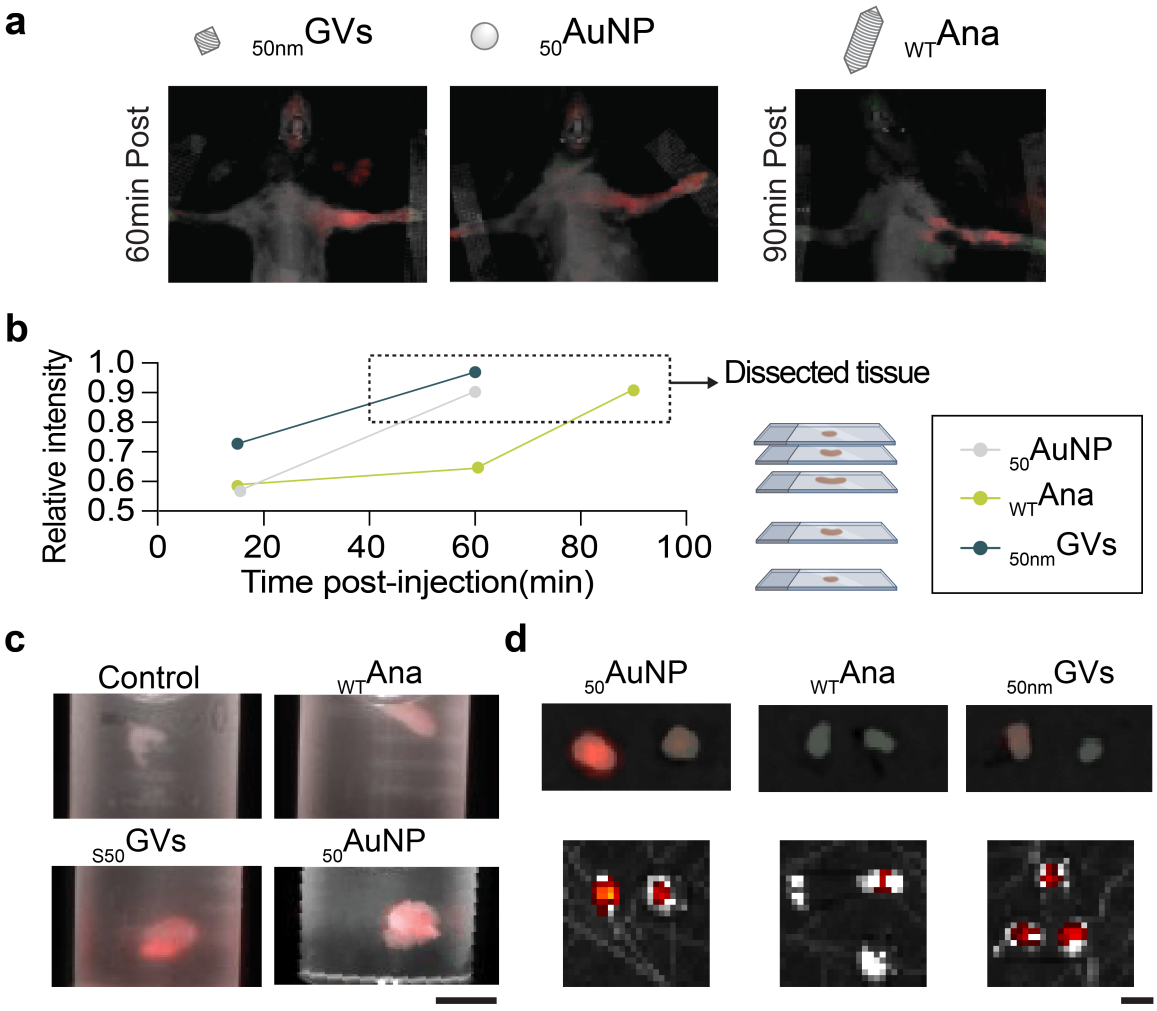


**Figure S2. Supplementary datasets illustrating the biodistribution of nanoparticles in lymph nodes.** a) A series of IVIS images showing the transportation kinetics of injected particles. Nanoparticles were indicated by a red channel. b) Relative fluorescent intensity changes within the targeted lymph nodes area at different time points. Lymph tissues were dissected when the intensity within the lymph node area reached a similar level as the injected sites. c) Overlay of epi-white light and fluorescence images of dissected mouse lymph nodes in PBS solution. The presence of fluorescently labeled nanoparticles is shown by the red color. Scale bar = 1mm. d) Additional IVIS images of dissected lymph nodes. Scale bar = 1mm.

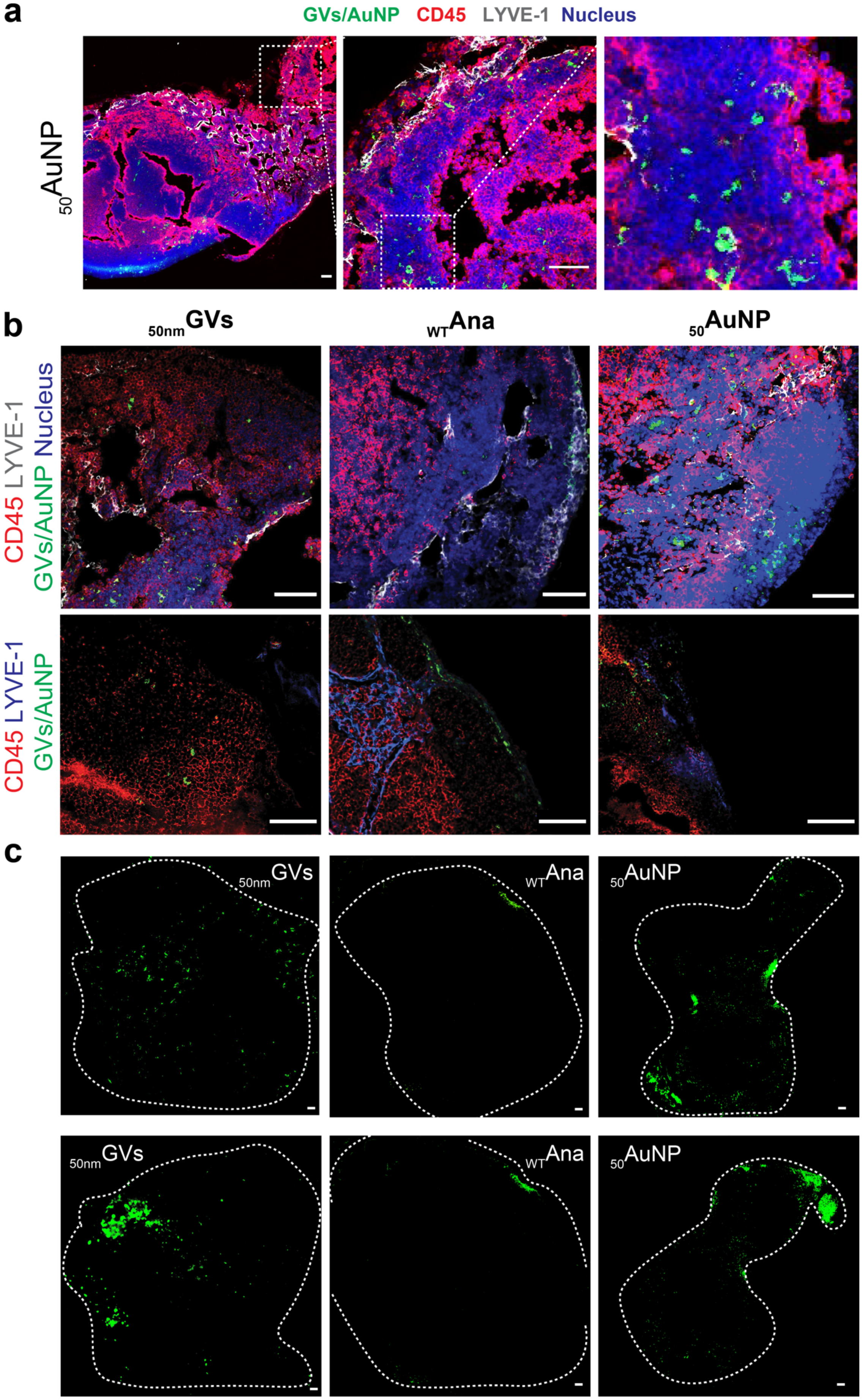
a
GVs/AuNP CD45 LYVE-1 Nucleus
50AuNP
b
50nmGVs
WTAna
50AuNP
CD45 LYVE-1 GVs/AuNP Nucleus
CD45 LYVE-1 GVs/AuNP
c
50nmGVs
WTAna
50AuNP

**Figure S3. Supplementary datasets illustrating the biodistribution of nanoparticles in lymph nodes.** a) Confocal fluorescence images of immunohistochemistry to depict the distribution of 50-nm gold nanoparticles ($_{50}$AuNP) within the lymph node tissue. The white dashed boxes in the left images outline the location of the areas zoomed in for the right images. Scale bar = 50 μm. b) Additional fluorescence images of immunohistochemistry were captured to depict the distribution of $_{WT}$Ana, $_{50nm}$GVs, and $_{50}$AuNP within the lymph node tissues. Each image is an overlay of multiple ones acquired in different fluorescent channels. The labeling on the left indicates the specific fluorescent channels used. Scale bars = 50 μm. c) Representative images demonstrating the distribution of nanoparticles inside the lymph node. The green fluorescence channel indicated the nanoparticles. The white dashed lines outline the periphery of lymph nodes on the slides. Scale bars = 50 μm.

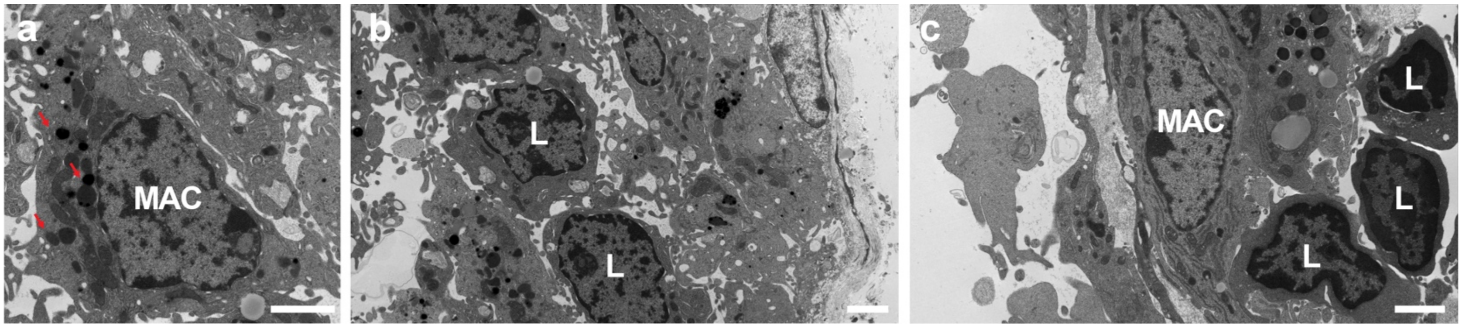


**Figure S4.** Representative TEM images of normal lymph node tissue. a) A representative TEM image of a normal macrophage. The typical features of a macrophage include a lobed nucleus and electron-dense lysosomes, indicated by red arrows. Scale bar = 2 μm. b) A representative TEM image of normal lymphocytes. Common lymphocytes exhibit a significant ratio of nucleus to cytoplasm. Scale bar = 2 μm. c) A representative TEM image of the edge area of the lymph node. Scale bar = 2 μm. The abbreviations used are MAC for macrophages and L for lymphocytes.
